# Local network-level integration mediates effects of transcranial Alternating Current Stimulation

**DOI:** 10.1101/216176

**Authors:** Marco Fuscà, Philipp Ruhnau, Toralf Neuling, Nathan Weisz

## Abstract

Transcranial alternating current stimulation (tACS) has been proposed as a tool to draw causal inferences on the role of oscillatory activity in cognitive functioning and has the potential to induce long-term changes in cerebral networks. However, the mechanisms of action of tACS are not yet clear, though previous modeling works have suggested that variability may be mediated by local and network-level brain states. We used magnetoencephalography (MEG) to record brain activity from 17 healthy participants as they kept their eyes open (EO) or closed (EC) while being stimulated either with sham, weak, or strong alpha-tACS using a montage commonly assumed to target occipital areas. We reconstructed the activity of sources in all stimulation conditions by means of beamforming. The analysis of resting-state data revealed an interaction of the external stimulation with the endogenous alpha power difference between EO and EC in the posterior cingulate. This region is remote from occipital cortex, which showed strongest EC vs. EO alpha modulation, thus suggesting state-dependency long-range effects of tACS. In a follow-up analysis of this online-tACS effect, we find evidence that this dependency effect could be mediated by functional network changes: connection strength from the precuneus, a region adjusting for a measure of network integration in the two states (EC vs. EO during no-tACS), was significantly correlated with the state-dependency effect in the posterior cingulate (during tACS). No analogous correlation could be found for alpha power modulations in occipital cortex. Altogether, this is the first strong evidence to illustrate how functional network architectures can shape tACS effects.

## Introduction

Local and long-range synchronized neural activity reflects coding and transfer of information in the brain, as studies have suggested over the years by *correlating* behavior with oscillatory components of electrophysiological recordings (Cohen & Kohn, 2011). In an attempt to infer more causal relationships between brain oscillations and cognition and behavior, diverse neurostimulation methods can be applied that manipulate neural activity. Among the non-invasive brain stimulation methods, transcranial alternating current stimulation (tACS) is gaining remarkable popularity. While online effects of transcranial electrical stimulation on behavior could be investigated, brain signals during stimulation were not recoverable due to its amplitude, several orders of magnitude higher than rhythmic neuronal activity. Yet, if the stimulation artifact is consistent and non-saturating, spatial or temporal filtering can recover brain signals, even during ongoing stimulation, as some research groups have been showing for the past few years (Soekadar et al., 2013; Helfrich et al., 2014b; see however Noury et al., 2016 and Neuling et al., 2017 for a rebuttal). Neuling et al. (2015) reconstructed brain activity from MEG recordings during simultaneously administered tACS. In this feasibility study, the authors resolved the well-known power increase in posterior alpha activity with eyes closed (Berger, 1929), showing that oscillatory effects can be recovered at the stimulation frequency. This innovation opens up the possibility to go beyond investigating *offline* aftereffects of the stimulation to understand *online* impact of tACS on brain dynamics.

State-dependent neural effects are a relevant trend established for other neurostimulation techniques (Neuling et al., 2013). This means that the cortex is more or less susceptible to the externally applied stimulation, depending on its fluctuating patterns of neural activity at local scales as well as on a network-level. State-dependency has broad implications: as the configuration of resting brain networks changes in many circumstances, most notably when damaged, treatment of neuropsychiatric disorders (Brittain et al., 2013) and non-clinical applications would benefit considerably from a clearer view of the nature of brain stimulation and its non-linear dependencies. Only a few studies have elucidated the online state-dependency of tACS. For example, a recent study by Alagapan et al. (2016) using modeling, invasive stimulation and ECoG in humans, showed that different behavioral states (such as task-engagement or resting EO / EC) profoundly impact how the same electrical stimulus perturbs ongoing activity. Non-invasively, alpha phase locking with the stimulation signal was shown to change as a function EO / EC state and current strength (Ruhnau et al., 2016b). No study so far has pursued the issue empirically as to what extent altered network states could mediate differential effects.

Effects of tACS are seen in brain regions distant from the stimulated cortex but anatomically and functionally connected (Cabal-Calderin et al., 2016). In a framework for explaining prestimulus predispositions on conscious near-threshold perception (Ruhnau et al., 2014), we argue that network-level integration of a neural ensemble determines its propensity to impact downstream regions. In practice, network integration has been operationalized via graph theoretical metrics (Frey et al., 2016), providing support for the framework. Here, we expect that an analogous mechanism could determine how tACS could exert long-range effects. Inter-hemispheric phase synchronization has been shown to be the mechanism behind tACS interference with functional coupling related to visual perception (Helfrich et al., 2014a). While the latter study shows tACS to affect brain connectivity, we ask how changes in functional network architectures could also shape the effect of tACS.

In the present study, we analyzed EO and EC resting-state MEG data from Neuling et al. (2015). We expected to see online increases of power at the stimulation frequency that are dependent on brain state and tACS. In testing the notion that altered functional network could have an influence on tACS effects, we took the most consistent inter-individual modulation of network integration (in EO vs. EC, in the precuneus) in the non-stimulated brain and investigated how this relates to the state-dependent alpha power change (in posterior cingulate) during tACS. This analysis revealed a significant association, whereas an analogous analysis using power changes during EO and EC (during no tACS) to correlate with the state-dependency effect was not significant. Overall, our results yield empirical evidence that functional network modulation mediated even by an apparently simple behavioural manipulation can profoundly shape tACS effects.

## Materials and methods

### Subjects

Seventeen healthy participants (9 males, 28 ± 4 years old; all right-handed) without psychiatric or neurological disorders volunteered for the study. The experiment was approved by the local ethics committee of the University of Trento and carried out in accordance with the Declaration of Helsinki. All participants gave written informed consent prior to its beginning.

### Stimuli and procedure

After applying the MEG coils and tACS electrodes to the head and determining the stimulation intensity (see next section), subjects were seated in an upright position in the MEG shielded room. After a brief resting-state measurement block and the estimation of the individual stimulation frequency (see *ISF determination* section below), participants were asked to keep their eyes open (EO) for 2 minutes until a tone and a visual instruction presented on a screen asked them to close their eyes (EC) for another 2 minutes. This was repeated three times while either no, weak, or strong tACS was applied (see below). The strong stimulation condition was always the last block, to avoid possible aftereffects. For more details on the complete experimental session, see Neuling et al. (2015).

### tACS parameters

A battery-operated stimulator system (DC-Stimulator Plus, NeuroConn GmbH, Ilmenau, Germany) was placed outside the magnetically shielded room. It was connected to the stimulation electrodes via the MRI module (NeuroConn GmbH). The stimulator delivered an alternating, sinusoidal current via two conductive-rubber electrodes (NeuroConn GmbH) of 7 by 5 cm applied with a conductive paste (Ten20, D.O. Weaver, Aurora, CO, USA) on the scalp at Cz and Oz of the international 10–20 system, chosen for maximal stimulation intensity in the parieto-occipital cortex (Neuling et al., 2012b). The electrode cables were located on the right side of the participant's head. In order to keep participants naive regarding the stimulation condition, the intensity was kept below the individual sensation and phosphene threshold (for each subject’s parameters, cf. Supplementary Table 1 in Neuling et al., 2015). To obtain the threshold, the subject was first familiarized with the skin sensation. Then the subject was stimulated with an intensity of 400 μA (peak-to-peak) at 10 Hz for 5 s. The intensity was increased by steps of 100 μA until the subject indicated skin sensation or phosphene perception or 1500 μA were reached. In the two cases in which the subject already reported an adverse effect at 400 μA, the intensity was reduced to a start level of 100 μA and increased by steps of 100 μA. The individual estimated threshold minus 100 μA was used as stimulation intensity in the strong tACS block. During the sham block, the experimental setup was the same as in the other blocks, but no electrical stimulation was applied. A stimulation intensity of 50 μA was delivered during the weak stimulation.

### MEG data recording

Electrophysiological brain activity was recorded at 1000 Hz (on-line band-pass hardware filters: 0.1-330 Hz) using whole head Elekta Neuromag MEG (Elekta Oy, Helsinki, Finland), housed in a magnetically shielded room (AK3b, Vacuumschmelze, Germany). Magnetic brain signal was spatially sampled at 102 positions, each consisting of a channel triplet of one magnetometer and two orthogonal planar gradiometers, yielding 306 sensors overall. Prior to the experiment, fiducials (nasion and left / right periauricular points), the location of five head position indicator (HPI) coils and more than 200 headshape samples were acquired for each participant with a Polhemus Fastrak digitizer (Polhemus, VT, USA). The coils tracked the position of the participants’ head during the experiment and the headshape served later for head modeling.

### ISF determination

Subjects’ alpha frequency, then used as the tACS frequency, was determined by analyzing the initial resting-state block immediately after the measurement (and before the stimulation blocks). The recording was divided into 2-second segments and for each we estimated the power spectra for frequencies ranging from 1 to 25 Hz in .25 Hz steps, after multiplication with a Hanning window and zero-padding of 4 seconds. Clear alpha peaks were identified in the averaged spectral power of the segments and of a chosen group of parieto-occipital gradiometers. In two participants, the peak was not evident from the data, so the frequency analysis was repeated for the EC portion of the first block (no-stimulation condition). The identified alpha peak was then used as the individual stimulation frequency (ISF, referred to as IAF in Neuling et al., 2015) for the tACS.

### Offline MEG data analysis

#### Preprocessing

Continuous data were offline high-pass filtered above 1 Hz and then downsampled to 512 Hz. Then EO and EC resting-state data were segmented into non-overlapping epochs of 2 s aligned to the phase of the stimulation. Epochs in the stimulation-free block were visually inspected to identify noisy, jumpy and dead sensors (then excluded from the whole data).

#### Source projection of raw data

Sensor space epochs were projected into source space using linearly constrained minimum variance beamformer filters (van Veen et al., 1997), following the procedure for single virtual sensors (www.fieldtriptoolbox.org/tutorial/shared/virtual_sensors) extended to 889 points covering the whole brain. These points were equally spaced by 1.5 cm in Montreal Neurological Institute (MNI) space and warped into individual head-space. The covariance matrix of each single trial was calculated and averaged across epoched data filtered from 1 to 40 Hz and then used to obtain the beamformer filters, together with single-shell head models (Nolte, 2003) derived from the individual head shape and the lead field matrix. Individual trials for each source location was obtained by multiplying the spatial filters with the sensor-level time series.

Beamforming is an adaptive procedure that optimizes spatial filters to restrict covariant signal from surface recordings to a source, localizing and spatially separating activity in the brain. This means that beamformer filters silence sources of noise correlated across sensors and suppress external artifacts in every location (Brookes et al., 2008). In addition to localizing activity in the brain, the aforementioned features seem tailored to reduce the tACS artifact, which is highly correlated noise. No regularization was used to avoid leakage of the tACS artifact to virtual sensors (Neuling et al., 2017). Of the 889 reconstructed sources, only the ones positioned strictly inside the MNI brain were used for subsequent statistical analysis, resulting in 576 virtual sensors. This was done to ensure that no peripheral source capturing the tACS stimulation was included in the statistics (Neuling et al., 2017). An MNI template brain was used for visualization purposes.

#### Resting-state power spectrum and PLV

Fourier coefficients were estimated for each epoch of the resting-state data in MEG sensors and in reconstructed activity in brain sources. We used a multitaper spectral estimation (Mitra and Pesaran, 1999) with a fixed smoothing window of ±2 Hz, with a 1 Hz resolution for the frequencies between 1 and 40 Hz and 2 Hz for those between 42 and 84 Hz. The same parameters for low and high frequencies bands were chosen in order to make comparisons across the entire spectrum.

Mean power densities were determined by averaging the squared absolute value (complex magnitude) of Fourier spectra across epochs and tapers for every sensor and brain source. As a measure of entrainment, we used inter-trial phase locking value (PLV; Lachaux et al., 1999), calculated as the absolute value of the mean of complex Fourier coefficients. As trials from sensor data were aligned to the phase of the tACS, this gave us an inter-trial coherence to the stimulation at the sources. These averages were computed separately for all stimulation conditions (sham, weak and strong tACS) and for the whole recording with both EO and EC states collapsed together or taken singularly.

#### Statistical analysis and state-dependency

Power from different conditions was compared using nonparametric cluster-based permutation dependent-samples T-statistics (Maris & Oostenveld, 2007). Cluster randomization was repeated over 5000 permutations. We contrasted relevant conditions at frequencies of interest, alpha and its first 2 harmonics, the first sub-harmonic and 2 control frequencies in between, corresponding to 5, 10, 15, 20, 25 and 30 Hz. Only cluster-corrected p values lower than 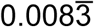 (0.05 alpha divided for the 6 frequencies comparisons) were taken into account. Dependency of tACS effects on EC and EO was assessed by computing the percent power change of strong tACS relative to no stimulation baseline within subject and condition. The normalized contrast for strong tACS state-dependency with EC and EO was computed with percent power change from no stimulation baseline within subject and condition, as follows:

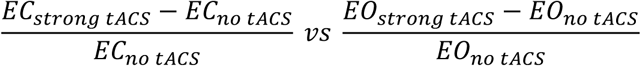

The same contrast was repeated for PLV and for weak stimulation. Because separate beamforming filters were used, it is not possible to draw a clear interpretation from a direct comparison between different levels of tACS on reconstructed brain sources, but this normalized contrast overcomes this limitation. For illustrative purposes, we also contrasted this way EC versus EO without stimulation and strong and no tACS conditions (both EC and EO).

#### Graph analysis

We calculated the source-by-source coherence from the Fourier spectra. The absolute value of the imaginary part of coherency was used as the functional connectivity metric to obtain the adjacency matrix. Imaginary coherency guards against spurious correlations due to volume conduction (Nolte et al., 2004). To find the best threshold to binarize the adjacency matrix, we started with the minimum value that ensured for each frequency the highest imaginary coherency, without disconnected nodes in the graph. This threshold, in which every node in the graph has at least one edge, avoided underestimation of connections. A range of thresholds around this value (±0.1 in steps of 0.02) was then tested to ensure the stability of the effects across choices of threshold. The threshold used was the one that had the highest and most stable effect for nearby values.

Local connectivity for each node was assessed with the following measures: node degree, efficiency, clustering and betweenness (Rubinov & Sporns, 2010). We then applied the same nonparametric cluster-based permutation tests to these measures as we did for power (also the same frequencies, see *Statistical analysis and state-dependency* section above), contrasting however only EC against EO for no tACS condition. Via this analysis, we assess the individual propensity of network modulation due to this subtle behavioural change, yielding a stronger argument that potential long-range effects during tACS are mediated by network-level changes. Local efficiency was the only measure that showed a cluster at a p value lower than 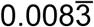. This index in particular, as the inverse of the graph average path lengths of all direct connections, reflects the integration of a node in the network. The source with maximum t-value was used as a seed to get individual imaginary coherency values to the source of maximum t-value for the state-dependency contrast. These values were used in the subsequently described seed-based partial correlations.

#### Regression and partial correlation

We wanted to explore whether the magnitude of the state dependency effect was correlated with connectivity. First, we assessed whether the different tACS intensities with which individual participants were stimulated would explain some of the variance of the effect (Ruhnau et al., 2016b). Therefore, we extracted for each participant the percent change of the strong tACS normalized EC / EO contrast (see *Statistical analysis and state-dependency* section) from the source with maximum group-level t-value. We then regressed the individual tACS intensity with these values. The linear model fit provided the adjusted R-squared and the F statistics.

As we ascertained the relation of stimulation intensity with state-dependency, we had to partial out the variance explained by intensity from subsequent correlations. We were interested in the area with differential network integration profile in EC and EO and its connection with the state-dependency region. We therefore took for each participant the EC / EO percent change of imaginary coherency from the source with maximal significant difference in local efficiency (see *Graph analysis* section above) to the state-dependency source. We then calculated the Spearman rank partial correlation coefficients between these individual percent changes of seeded imaginary coherency and state-dependency, controlling for tACS intensity. We repeated the partial correlation also with percent change of EC / EO power in the occipital cortex instead of seeded coherency, also to control for spurious correlation due to the most variance explained by tACS intensity. Correlation coefficients were t-tested against the two-sided alternative hypothesis of no partial correlation.

## Results

### Power modulations during tACS

The cluster-based statistic on 10Hz contrasting the strong tACS against sham (calculated with separate leadfield filters) portrayed a strong increase for the stimulation condition distributed in the whole brain, maximally localized in the border between the cuneus and the precuneus (*Figure 1a*). The difference was highly significant (p_cluster_ < 0.001), but its interpretation is unclear (see *Discussion*) and it is shown here only for illustrative purposes.

**Figure 1.**
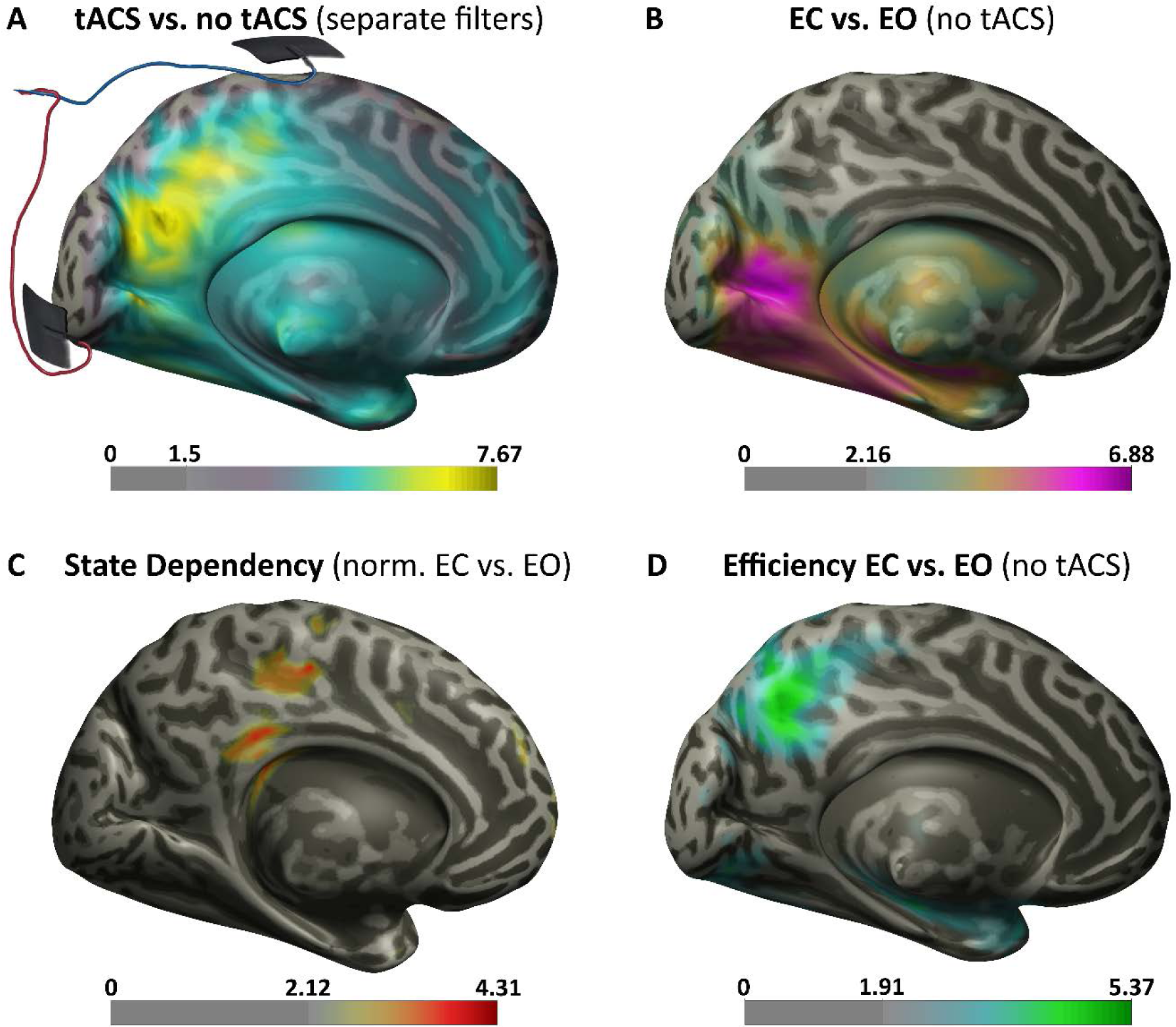
Posterior t-value Topographies. Significant t-values brain maps of the principal effects at 10 Hz during EC or EO and no or strong alpha tACS. On the bottom of every MNI hemisphere, the color bars with the t-values masked for cluster-corrected p < 0.05 significance. Between 0 and the max is the lowest significant t-value. **A.** Medial view of the left hemisphere showing the contrast between strong and no tACS. Wired rubber patches show the location of the electrodes (attached to the scalp) with which the stimulation was delivered. The strongest effect is located in the medial parieto-occipital cortex, but an interpretation of this contrast is risky (see text). **B.** The well-known increase in alpha power in EC contrasted with EO in occipital and parietal regions. All the tACS conditions showed this difference, but here only the no stimulation contrast is shown. Alpha power differences in this region were used as predictors in the partial correlation of *Figure 2b.* **C.** A stronger alpha increase for EC relative to EO during stimulation in the posterior cingulate, obtained with a normalized contrast denoting state-dependency. The max t-value source was the target of the correlations and seeded coherency of *Figure 2* and *Figure S3*. Unlike the other brains in this figure, here the right hemisphere instead of the left is shown. **D.** Local efficiency increase in EC relative to EO in the precuneus during no tACS. The source with the strongest contrast was the seed of imaginary coherency used in the partial correlation of *Figure 2a.*

As expected, power at 10 Hz increased in contrasts between EC and EO for all tested frequencies in all stimulation conditions (p_cluster_ < 0.002 for no stimulation, *Figure 1b*), but there was no PLV change across all conditions (*Figure S2c*). The difference was maximally localized in parieto-occipital areas. Again, this well-established pattern is shown here only to illustrate the topography and its relation with the other effects.

Ongoing brain oscillatory activity interactions of stimulation and brain state was assessed with a stimulation-normalized contrast between EC and EO. No state-dependent power activity was found for this contrast for weak tACS. For strong stimulation, the cluster-based permutation test revealed a significant difference (p_cluster_ < 0.006) at 10 Hz and the effect was maximally expressed in the posterior cingulate (*Figure 1c*). This indicates that this is the area where state-dependency at the alpha frequency was maximal, that is where the power enhancement driven by the external stimulation was modulated also by the EC brain state. The effect was only observed for power, and no significant interaction nor change in PLV was found (*Figure S1b* and *Figure S1c)*.

At 30 Hz, the state-dependency contrast exposed a difference in power (p_cluster_ < 0.002), most pronounced in the right superior frontal gyrus (*Figure S1d*). This online cross-frequency modulation related to EO / EC brain state was again driven by power resonance (*Figure S1e*) but not by inter-trial PLV (*Figure S1f*). As in the case of alpha, we cannot assert anything about the sharp increase in frontal 30 Hz power and PLV in strong tACS relative to sham stimulation, because we cannot compare conditions. No other tested frequency showed state-dependent effects.

As these previous analyses established a state-dependent modulation remote to regions showing the primary power change between EC and EO (i.e. posterior cingulate vs. occipital cortex), this implied differences in neural states between these two conditions to mediate this long-range effect. This is the issue pursued in the next section.

### Network-level changes between EC and EO

To expose differences in brain network states between EC and EO, which could potentially influence this novel state-dependent effect, we used graph theoretical analysis of the resting data without tACS. The whole-brain nonparametric statistic on local efficiency uncovered a cluster in the precuneus for the 10Hz band, which showed a significant increase in EC relative to EO (p_cluster_ < 0.004, *Figure 1d*). Unlike the other frequency bands and the other graph measures, the effect was stable across a range of values for adjacency matrix thresholding, between 0.12 and 0.26 of absolute imaginary coherency, with a maximum effect at 0.18.

### Effects mediating state-dependency

As participants received different tACS strength, we examined the correlation between stimulation intensity and the normalized difference in power denoting state-dependency. We found a correlation between individual intensity and the state-dependent effect (adjusted R^2^ = 0.24, p = 0.027; *Figure S3*).

To characterize the relationship of the underlying pattern of connectivity with the state-dependency, the source with maximum statistical value in the efficiency contrast (precuneus, *Figure 1d*) was used as a seed to extract imaginary coherency to the region of the cortex with the state-dependent effect, the posterior cingulate (*Figure 1c*). The partial correlation controlling for tACS intensity revealed that the connection strength between precuneus and posterior cingulate was a predictor of state-dependency (Spearman rho = 0.53, p = 0.036; *Figure 2a*). Still controlling for intensity but using EC / EO alpha power difference as the predicting variable (extracted from the source in the occipital cortex with the strongest contrast; *Figure 1b*) showed no correlation with state-dependency (Spearman rho = −0.24, p = 0.35; *Figure 2b*).

**Figure 2.**
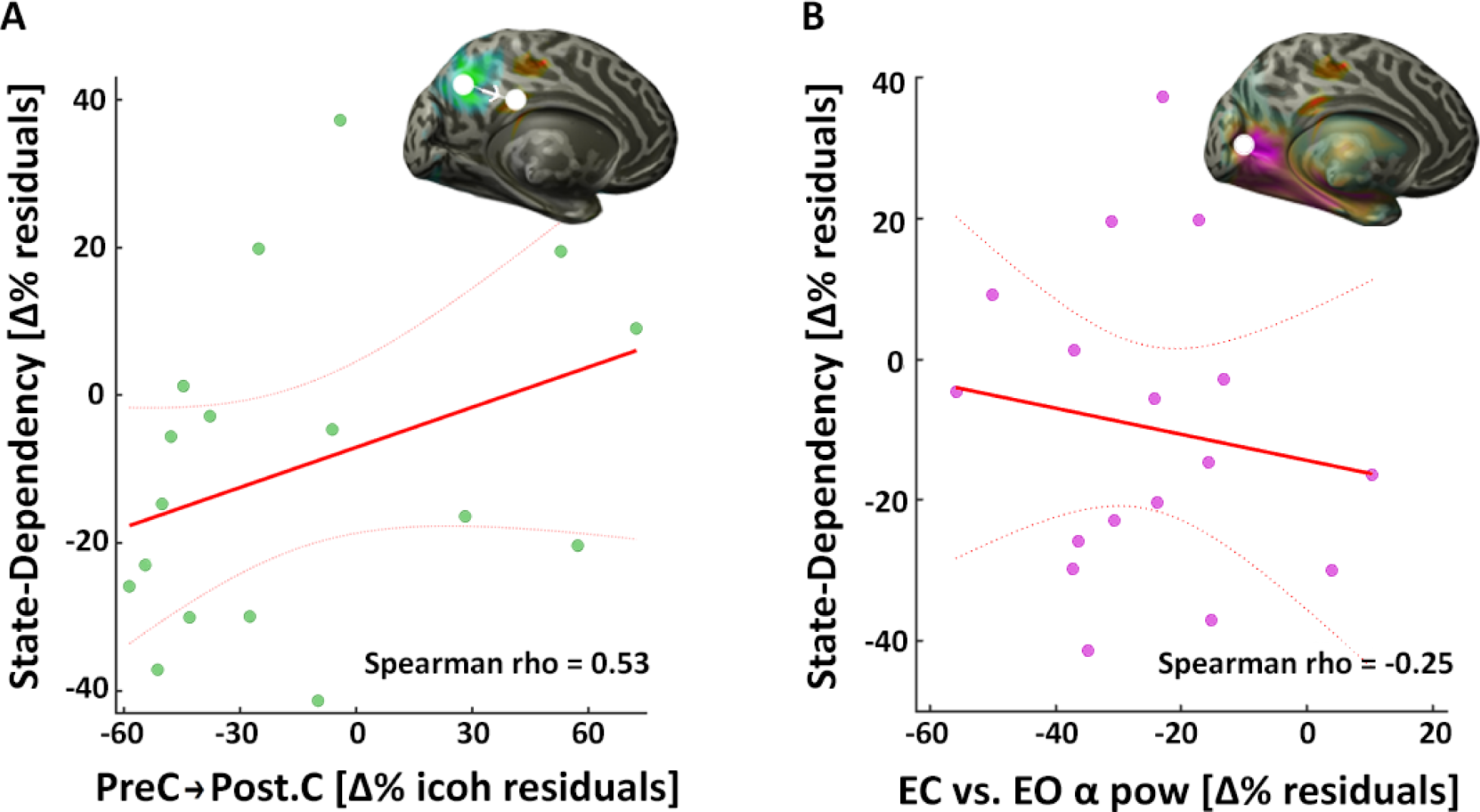
Partial Correlations. Correlations and linear model fits to the individual state-dependency percentage changes. Dotted lines represent the 95% confidence intervals. **A.** Linear model fit of the adjusted values (residuals) of the partial correlation between imaginary coherency (icoh) from the source with maximum local efficiency difference, the precuneus (PreC, *Figure 1d*), to the state-dependency one in the posterior cingulate (Post.C, *Figure 1c*, as the inset on the top right shows). The controlled variable is stimulation intensity, the independent variable in *Figure S1*, here partialed-out. Again, for both the predictor icoh and the predicted state-dependency, these are individual percent change of EC vs. EO. **B.** Same as in *A.*, but with alpha power increase in EC vs. EO as a predictor (extracted from the source with maximum t-value, as the inset and as *Figure 1b*). This correlation is non-significant.

## Discussion

### State-dependent power effect of tACS

Our analyses uncovered online interactions of tACS with brain state at the alpha frequency. Power enhancements driven by external stimulation interacted with endogenous alpha increases in the transition from EO to EC. The impact of tACS changed depending on brain state: it was more pronounced during EC in the posterior cingulate (*Figure 1c*), an area between the stimulation electrodes but where current flow is not at its peak (*Figure 1a*, but see below). This could be due to the dominant Eigenfrequency increase in posterior regions following closing the eyes, which overshadows the tACS influence (*Figure 1b*, seen also in offline aftereffects in Helfrich et al., 2014b). That is, neural firing rate in occipital and parietal regions might still be influenced by tACS differently in the EC brain state with respect to EO, but if present, this effect is too small relative to the endogenous power modulation. On the other hand, the posterior cingulate cortex was affected differently by the stimulation, with an unclear involvement of resting networks and with consequences for task-active cognition (Leech et al., 2013). This speaks for the specificity of tACS efficacy, which is seen in certain tasks, setups and sensory modalities but not in others. The fact that it is part of the default mode circuit (with distinctive tACS network effects; Cabal-Calderin et al., 2016) and its relation with graph measures (discussed below) reinforce the idea of the singularity of this region.

When contrasting induced alpha power for different tACS conditions, there were differences distributed across the whole brain (*Figure 1a*). This is difficult to interpret, as the variance was probably also driven by the different beamformer filters obtained separately from different conditions and their interaction with the reconstructed alpha power. As the filters have to suppress the artefact in the strong tACS condition, their profile is very different from the ones during no stimulation. However, one can appreciate how the cluster reflects maximum current flow of tACS located between the electrodes, as it has been modeled (Neuling et al., 2012b). Even though there is no information on tissue density in our analysis, the effect in source reconstruction shows the current diffusion where the finite-elements model predicts it.

The state-dependent effect of tACS at 30 Hz showed a frontal source distribution and again there was a higher increase in the induced power in the EC state (*Figure S1d-e*). Interconnected areas respond, even if not directly stimulated, usually in the range of the natural frequency of their local cortico-thalamic circuit (Rosanova et al., 2009). This cluster in the right frontal pole in the second harmonic (30 Hz) follows this preferred pattern. This state-dependent interaction is reminiscent of the cross-frequency impact that tACS has on the brain, especially on multiples of the stimulation frequency (Neuling et al., 2017). It is hard to declare what drives this modulation, be it EO / EC differential frontal activity (Barry et al., 2007), resting-state network connections from the posterior cingulate, excitability for the preferred frequency of the region (Rosanova et al., 2009), or an interaction between these factors during stimulation. The uncertain nature of this effect at 30 Hz prompted us to stop enquiring further. Yet again, harmonics and their brain state dependencies are another variable to consider when applying tACS.

### Effects of tACS not dependent on simple entrainment

The interaction of tACS with EC seems to be driven only by power resonance, without a change in PLV (*Figure S1b*). The posterior topography of higher alpha in EC relative to EO, seen in all tACS conditions as clearly as during no stimulation (*Figure 2b*), was likewise not accompanied by phase alignment. This means that a strong continuous entrainment, in which endogenous oscillators align their phase and their frequency to the stimulation, is absent and does not contribute to power increase or online state-dependency of tACS. A trend of reduction in synchrony measured by PLV was present in the posterior cingulate but not in the parieto-occipital areas. This state-dependence of the tACS on the well-known basic alpha dynamic, adjusting when eyes are closed, is therefore unrelated with phase synchrony across trials.

This may be surprising (Herrmann et al., 2013), but there is some debate on whether entrainment is really the mechanism behind tACS effects. Many tACS studies show phasic modulation of behavior (e.g. Neuling et al., 2012a), which can be attributed to entrainment. State-dependent stimulation-driven phase alignment can be seen after trial averaging (Ruhnau et al., 2016b). Conversely, in another online tACS study, effects on PLV (same formula used in this study) are missing, if not even dampened, at the simulation frequency (Ruhnau et al., 2016a), even though there were several oscillators driven at the same time.

Inter-trial phase coherence is not associated to the increase in source power, especially in this tACS setup (Neuling et al., 2013). Rather than an instantaneous synchronization of endogenous oscillators, tACS has been proposed to be altering synaptic strength, as its mechanism of action to modify brain frequencies (Vossen et al., 2014). Alternate current stimulation effects related to phase are also not seen also in recordings with implanted electrodes (Opitz et al., 2016; Lafon et al., 2017; but see Noury et al., 2017, also for a reproach on phase estimates in deep-brain recordings).

### Brain connectivity behind state-dependency

Functional connectivity fluctuates continuously, but it varies mostly when the brain changes its *mode* of functioning. This happens during changes between different tasks or the diverse repertoire of states like vigilance or attention. Even a simple change like closing the eyes creates profound reorganizations of brain activity. In this case, parietal cortices exhibit modifications in their oscillatory balance, especially in the low frequency bands, closely accompanied by changes in connectivity in the same frequencies. In the alpha frequency, the most stable change seems to be the integration of the parietal cortex with the rest of the brain, which increases when eyes are closed (*Figure 1d*). It is reasonable to assume that the variation in connectivity from this region to the rest of the brain is bound to influence other processes that ride on these functional connections (Ruhnau et al., 2014), such as activity modulations caused by tACS.

Reasonably, it appears that tACS influence dependent on ongoing brain state is mostly driven by the intensity of the electrical stimulation (*Figure S3*). On top of that, regional variation in the system’s connectivity are related to the amount of this state-dependency. Namely, the connection strength increase to this region is followed by a better integration and efficiency (*Figure 2a*). Regional alpha power modulations as commonly seen during EC vs. EO, however, do not seem to mediate this state-dependent effect of tACS (*Figure 2b*).

Since functional connectivity indicates electrical information flow among brain areas, it would be evocative to think that they also control the resistance of tACS diffusivity. The current flow at the neuronal scale should be the same across conditions (*Figure 1a*) and changes in functional connections during different brain states reflect the distinctive responses to the impact of tACS. In other words, connectivity controls and follows patterns of inhibition and excitation in oscillatory activity, so it is likely to interact with tACS. Crucially, this is revealed in cortical regions distinct from the maximum tACS induced current.

## Conclusion

As non-invasive electrical brain stimulation, particularly tACS, becomes more and more popular, caveats are bound to emerge. Although there is some convincing evidence on the reliability of tACS and its impact on behavior, its direct influence on brain processes is unknown. For one thing, the present consensus was that tACS works by entrainment and that an online alignment of endogenous oscillation would follow. Instead, in this study, we have seen how during stimulation the phase of the stimulated frequency is not impacted in this setup, and rather, tACS seems to interfere with it across trials. If a sub-neural ensemble is entrained, it is not large enough to be the dominant pattern in inter-trial PLV, but there is yet no strong support on the current situation at this level.

The main message of this study is that the online effects of tACS in the parenchyma is also defined by functional factors. In the case of tACS, oscillatory activity and connectivity are interwoven with both its ability to affect interconnected regions that are far from the focal stimulated zone and its basic impact on the cortex. As we have seen, a modulation of the functional network architecture by the simplest possible behavioural manipulation has a profound impact on tACS effects. The absence of correlation for power shows that the effects are specifically mediated via network reorganization. This finding yields important confirmation that it is connectedness that matters in determining whether activity in a region will affect downstream areas (Ruhnau et al., 2014). As modeling and predicting effects become more complex, this should have profound implications for the use of tACS in cognitive and clinical neuroscience, i.e. one should be wary when treating patients based on results obtained in controls.

Researchers must control many variables in a neurostimulatory experiment, in which varying a single parameter can reverse its effect (Moliadze et al., 2012; Veniero et al., 2017). Alternate current stimulation is alluring: if it can manipulate physiological frequencies directly, then we can assess whether oscillatory activity actually plays a causal role in cognition and behavior. However, the state-dependency of tACS effects, as well as their nonlinearity, make these current causality assumptions naïve and inadequate. Thus, it is important to take this into account, both for researchers, applying tasks with distinctive activity and connectivity profiles and clinicians, who often have to deal with brain systems that have altered network organizations.

## Acknowledgements

This project was supported by the European Research Council (ERC StG 283404). We thank Dr. Elie El Rassi and Dr. Benjamin Timberlake for comments on an earlier version of the manuscript.

---

**Figure S1.**
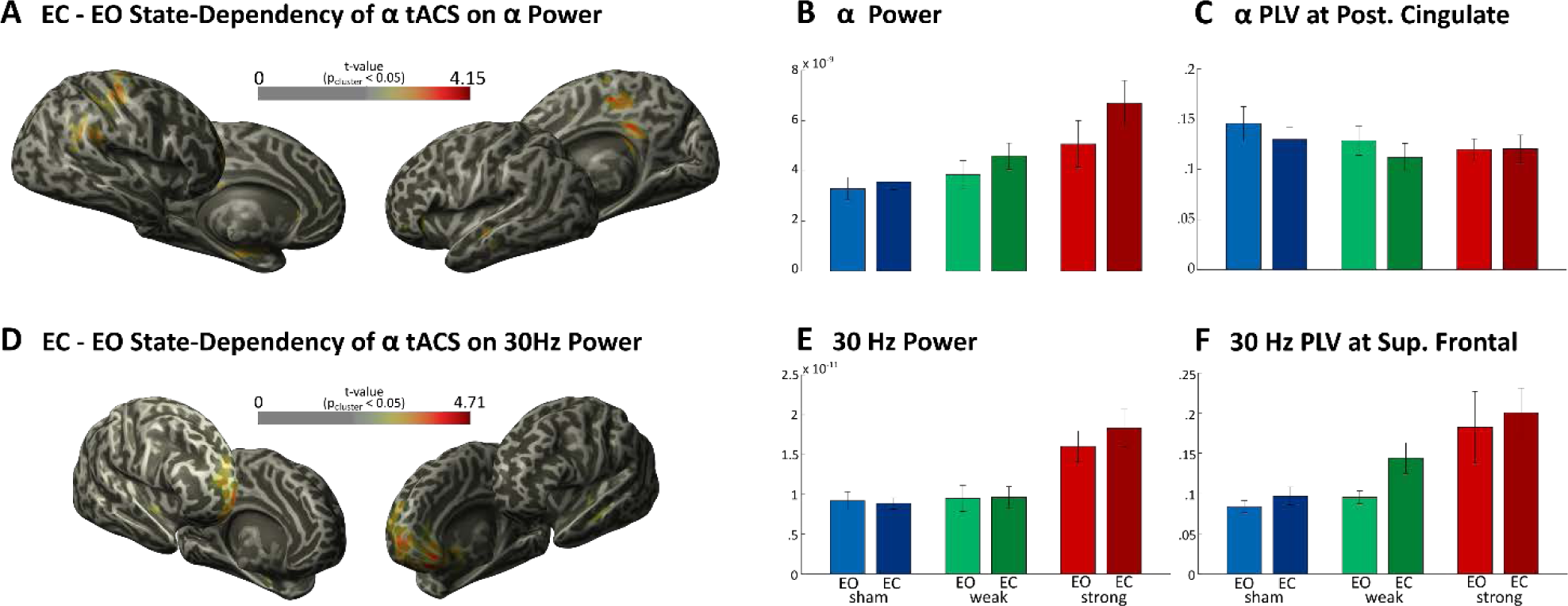
State-Dependent Effects. Strong alpha tACS and EC / EO state interactions at 10 Hz and at 30 Hz. **A.** The projection on the left and right views of the left and right MNI hemispheres (left -medial-view of right hemisphere on the top right is the rotated *Figure 1c*) of the contrast exposing the topography of a stronger alpha increase for EC relative to EO during stimulation in the posterior cingulate. On the top, the color bar with the t-values masked for cluster-corrected p < 0.05 significance. **B.** Alpha Power in the virtual sensor with maximum t-value in the posterior cingulate (MNI cords [10 −35 25] mm, target of the correlations and seeded coherency, *Figure 2*) at different stimulation and EO / EC conditions. The state-dependent effect in power capture by the normalization is discernible from the bigger EO / EC change in strong tACS. **C.** Alpha PLV at the same source. Power increase and the slight PLV decrease visible in the progressive levels of stimulation could be due to filter difference as stated before, so we cannot confidently state anything about that. **D.** Same as in *A*, but the brain is rotated to better show the EO / EC state-dependency at 30 Hz of alpha tACS in the Superior Frontal Gyrus. **E.** Power at 30 Hz in the Superior Frontal Gyrus source (MNI cords [25 70 −5] mm) in all conditions. **F.** PLV at 30Hz at the same position. Bars indicate the condition mean and errorbars the within-subjects 95% confidence intervals.

**Figure S2.**
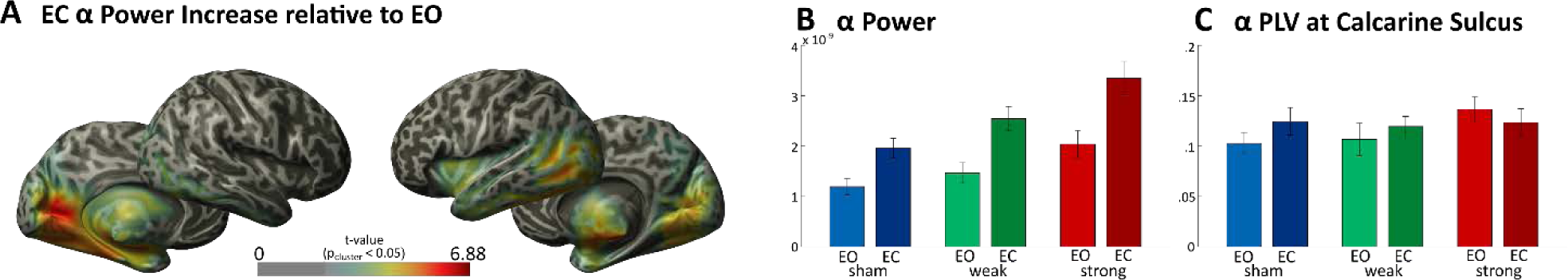
EC Alpha Power Increase. Higher power in EC state relative to EO in occipital regions. **A.** Same as in *Figure S1*, but with the posterior topography of higher alpha power in EC relative to EO (right -medial-view of left hemisphere on the bottom left is the same as *Figure 1b*). **B.** Alpha Power at the Calcarine Sulcus (MNI cords [−5 −95 −5] mm, max t-value, one of the seeds for coherency used in the partial correlations, *Figure 2b*) for the different conditions. **C.** Alpha PLV at the same location. Bars indicate the condition mean and errorbars the within-subjects 95% confidence intervals.

**Figure S3.**
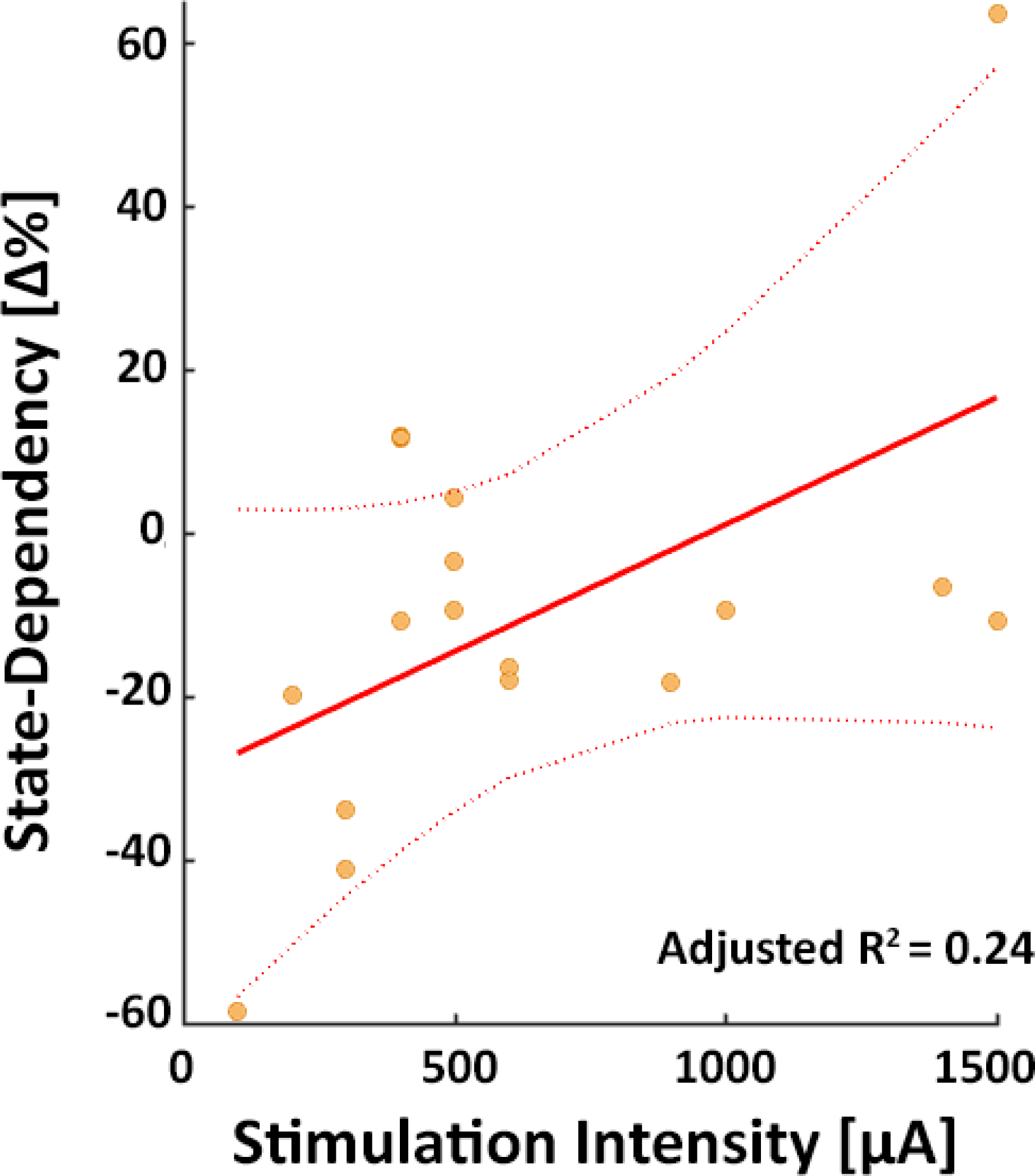
Regression between Intensity and State-Dependency. Correlation between the strong tACS intensity and values for every participant of the normalized change used to detect state-dependency, individual percent change. Dotted lines represent the 95% confidence intervals.

